# PARP1 Regulates 3D Structure and Function of Super-Enhancers and Hormone-Control Regions in Breast Cancer Cells

**DOI:** 10.1101/2025.08.01.668155

**Authors:** Houda Baccara, Francois Le Dily, Davide Carnevali, Jofre Font-Mateu, Cristina Moreta-Moraleda, Alex Franco Galvez, José Carbonell-Caballero, Martin Teichmann, Miguel Beato, Roni H. G. Wright, Roberto Ferrari

## Abstract

Poly (ADP-ribose) polymerase 1 (PARP1) has been linked to various genomic pathways and in the maintenance of genomic stability. PARP1 has also been postulated to regulate cis-regulatory elements, but the mechanisms remain largely unknown. Here, we employed genomic approaches to show how PARP1 occupies Super-Enhancers (SEs) of luminal A T47D breast cancer cells and how some PARP1 is dynamically relodged by progesterone at hormone-control regions (HCRs). Disruption of PARP1 causes transcriptome reprogramming, altering the expression of both SE-associated and HCRs-regulated genes. This is achieved through a dual coordinated action in the establishment of long-range and intra-TADs chromatin loops, which are lost upon PARP1 ablation, but enforced by its catalytic inhibition. We show that PARP1 also plays a role in regulation of chromatin compartments. Our results reveal PARP1, as a new chromatin looper and compartments organizer controlling SEs and HCRs genes linked to cell identity and hormonal response in breast cancer cells via 3D genome organization.

**Highlights (Separate document):**

- PARP1 occupies cell specific super-enhancers and hormone-control regions.
- Loss of PARP1 alters the cell identity and progestin-induced gene expression signature.
- Gene expression changes upon PARP1 loss relay on altered 3D chromatin looping and compartments.

**eTOC blurb:** We have discovered that the nuclear repair protein and cancer drug target; PARP1 is preferentially bound at super-enhancers. The binding of PARP1 is essential for the underlying, cell specific gene expression program and the 3D structure of the chromatin surrounding the super-enhancer regions.

**Graphical abstract:** Separate document

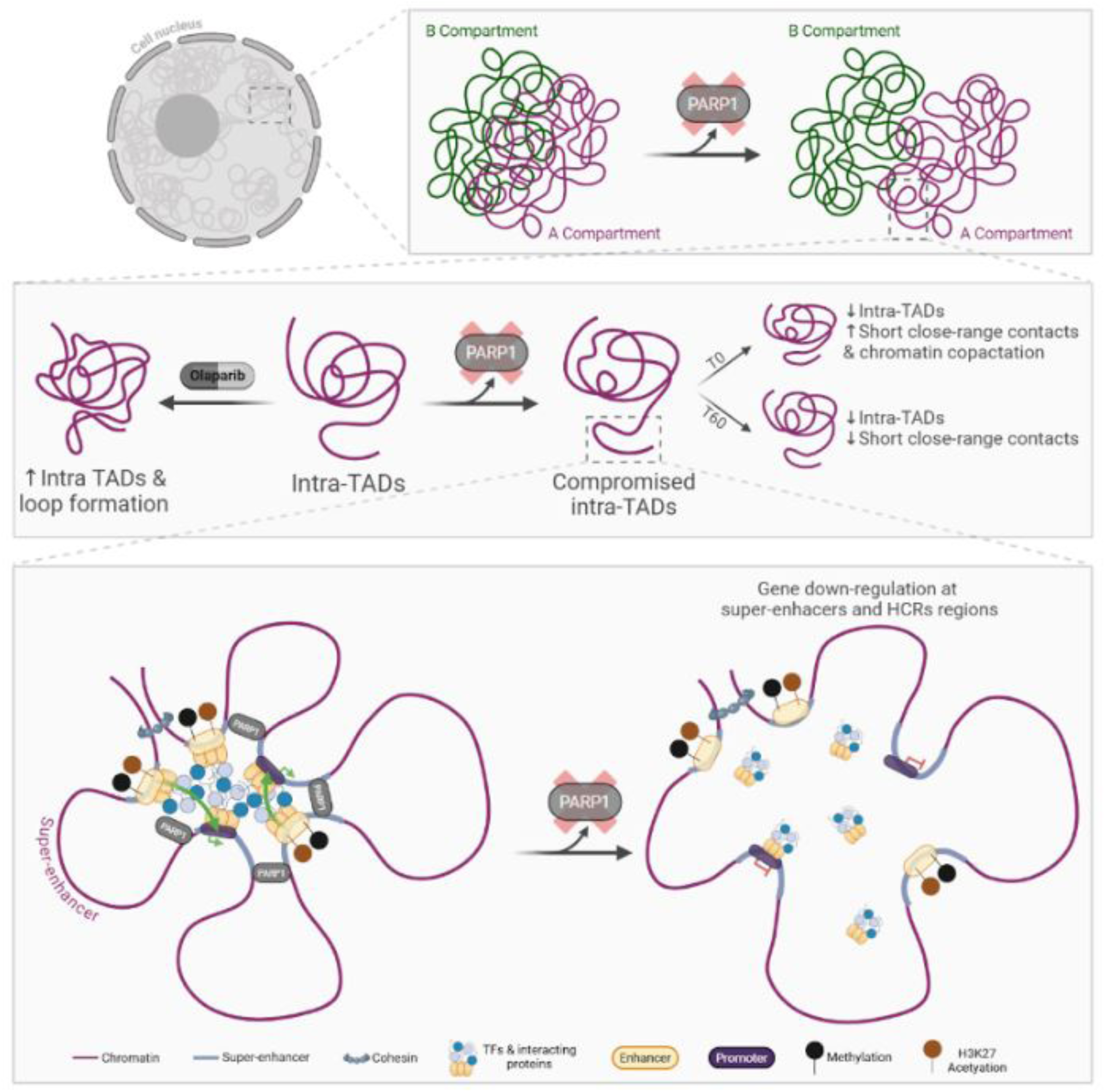

## Introduction

Poly(ADP-ribose) polymerase 1 (PARP1) is a multifaceted enzyme known primarily for its involvement in DNA repair processes (Huang and Kraus, 2022). However, beyond its well-established roles in the maintenance of genomic integrity, PARP1 has emerged as a pivotal player in the regulation of gene expression (Huang and Kraus, 2022) and in orchestrating epigenetic events setting up chromatin domains (Ciccarone et al., 2017). PARP1 catalyzes the transfer of ADP-ribose units from NAD^+^ to target proteins, thereby modulating chromatin structure and accessibility (Alemasova and Lavrik, 2019). This modification, known as PARylation, plays a critical role in the transcriptional regulation by altering the binding dynamics of transcription factors and promoting or inhibiting gene transcription depending on the context (Bordet et al., 2020), for example controlling progesterone action (Wright and Beato, 2012, Wright et al., 2012). Moreover, PARP1’s influence extends to genome architectural roles, albeit its involvement in 3D genome structure was only proven in the context of Epstein-Barr virus (EBV) latency via PARP1 interaction with chromatin architectural proteins CCCTC-binding factor (CTCF) and cohesion (Morgan et al., 2022). PARP1 catalytic inhibition altered the EBV episome’s 3D structure, affecting gene expression at specific chromatin looping regions (Morgan et al., 2022). Nevertheless, in human, PARP1 role in 3D genome organization remains elusive due to the lack of high-quality mapping of its genomewide occupancy. Here, through successful ChIP-sequencing of PARP1 in breast cancer cells, we unveiled its occupancy and ability to establish long-range chromatin loops of Super-Enhancers (SEs) and Hormone-Control Regions (HCRs) with their regulated genes, independently of CTCF and its PARylation activity. Our data provide evidence of a 3D architectural looping role of PARP1 in human cells which also extends to the regulation of chromatin compartments.

## Results

### PARP1 binds Super-Enhancers and is dynamically redistributed upon progestin exposure to Hormone-Control Regions in breast cancer cells

So far PARP1 ChIP-seq attempts failed to report significant enrichment at specific genomic sites and therefore the precise chromatin occupancy of PARP1 (Figure S1A). Here we combined a tested ChIP-grade antibody (AB_2793715, Active Motif) and High-sensitivity ChIP protocol (ChIP-IT High Sensitivity®, Active Motif) to identify the genomewide occupancy of PARP1 in T47D breast cancer cells before and after progestin (R5020) treatment. Our approach yielded high quality mapping of PARP1 revealing 13999 peaks before (T0 condition) and 17094 after 1h hormone exposure (T60 condition) (Figure 1A). The two sets of peaks showed a high degree of overlaps with 9226 common regions (Figure 1A). Compared to previous PARP1 ChIP-seq experiments from HEK293T cells (GSM1938987) and MCF-7 cells (Nalabothula et al., 2015), (Figure S1A) our PARP1 ChIP-seq clearly showed precise peaks of enrichment with very little background noise (Figure S1A). To gain insight into the functional occupancy of PARP1-bound regions across our two conditions we employed combinatorial clustering of PARP1 peaks (Figure 1A) and plotted the heat map of the three different classes of PARP1-bound genomic regions (Figure 1B). The first cluster (named T0) showed significant enrichment for PARP1 only at T0. The second (named common) revealed unchanged levels of PARP1 occupancy in both conditions, and the third cluster (named T60) was enriched for PARP1 only upon hormone treatment of 60 min (Figure 1B). To gain insight into the functional occupancy of these PAPR1-bound regions we interrogated Cistrome Toolkit (Zheng et al., 2019) to identify putative chromatin regulators associated with the three clusters (Figure 1C). We found that PARP1 occupancy at clusters T0 and common was strongly overlapping with several chromatin proteins enriched to super-enhancers (SEs) (large clusters of typical enhancers driving expression of cell identity genes) (Hnisz et al., 2013). Among these we find Bromodomain 4 (BRD4), Mediator 1 (MED1) and RNA polymerase II (POL2) (Figure 1C, upper panel), suggesting a putative broader role at cis-regulatory regions dependent on PGR. It is known that PGR can cluster in specific genomic locations driving progesterone target gene expression reprogramming, referred to as T47D Hormone-Control Regions (T47D-HCRs) (Le Dily et al., 2019b) The cluster analysis of T60 PARP1 binding sites (Figure 1C, lower panel), led us to hypothesize that PARP1 could also be enriched at these T47D-HCRs together with SEs. First, to prove PARP1 is enriched at SEs, we collected T47D SEs reported in the SE-database (SEdB) (T47D-SEdB) (Wang et al., 2023) and searched for overlaps with PARP1 clusters. We found that about 90% of T47D-SEdB overlapped with cluster T0 and common (Figure 1D, upper panel). Similar analysis was performed using T47D-HCRs sites and we unveiled that 80% of overlap with cluster T60 (Figure1D, lower panel). Hence, if PARP1 is a bona fide SEs interactor it could be used to predict SEs in T47D similar to what has been described for H3K27ac (Whyte et al., 2013). In order to examine PARP1 as a SE regulator, we firstly calculated the Pearson correlations between our PARP1 ChIP-seq with several published T47D ChIP-seq datasets of active and repressive histone marks (H3K18ac, H3K27ac, H3K36me3, H3K27me3 and H3K9me3) and SE regulatory proteins (POL2, CTCF, PGR and BRD4) (Ferrari et al., 2020, Zaurin et al., 2021). Clustering the calculated correlation coefficients showed PARP1 was strongly correlated with H3K27ac (Sup. Figure S1B), attesting at its functional relationship with this active mark. We then used ChIP-seq of H3K27ac at time T0 and calculated de-novo T47D SEs using ROSE algorithm (Loven et al., 2013, Whyte et al., 2013) (Supp. Figure S1C) and compared it to T0 PARP1-calculated prediction (Figure 1E). Our results show how PARP1 was able to predict a large fraction (63%) of SEs calculated with H3K27ac (Sup. Figure 1D), and other new potential SEs that were not detected by histone acetylation but strongly enriched in PARP1 (Figure S1D). Analysis of the genes associated with PARP1-derived SEs (PARP1-SEs) showed a strong degree of overlap with H3K27ac-derived SEs (H3K27ac-SEs), with the strongest SEs associated with the same genes (Figure 1E and Figure S1C). We then compared PARP1-SEs, T47D-SEdb and a set of control SEs from Keratinocytes (KT-SEs) (from the SEdb) for enrichment of PARP1, BRD4 and POL2 at time T0 (Figure 1F). PARP1-SEs revealed far greater enrichment for all three marks compared to T47D-SEdb and KT-SEs, demonstrating that PARP1 chromatin enrichment alone is capable of better predicting SEs in a cell line specific manner (Figure 1F). Among all T47D-SEdb, one important SE was known to be associated to the GSE1 gene. Our analysis here shows that both PARP1 and H3K27ac ChIP-seq were capable of de-novo predicting this SE (Figure 1G), both exhibited strong overlapping enrichments at this locus together with BRD4 (Figure 1G).

**Figure 1.**
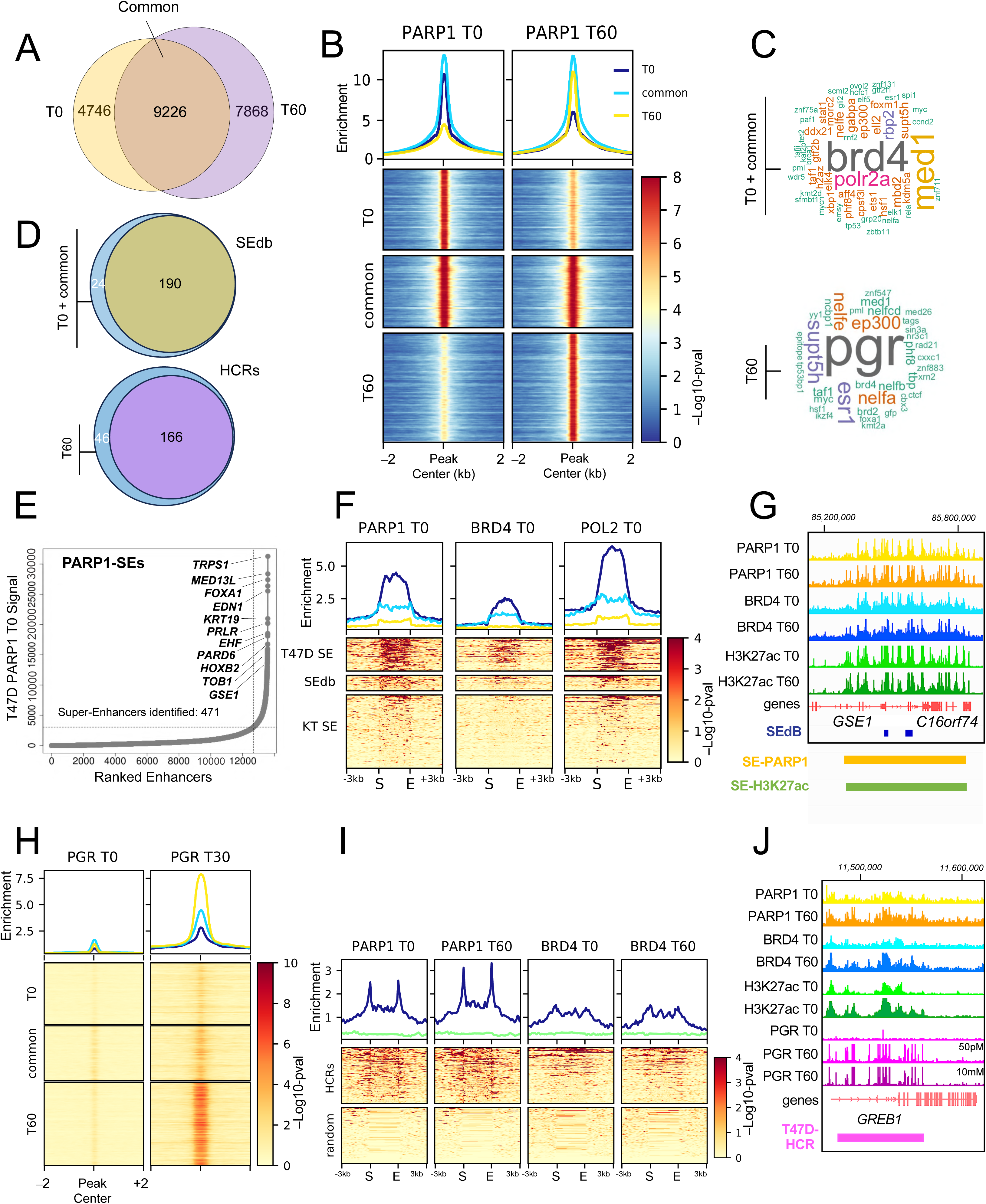
PARP1 genome wide occupancy of Super-Enhancers (SEs) and Hormone-Control Regions (HCRs) in T47D breast cancer cells. **A)** Venn diagram showing the overlap of PARP1 ChIP-seq peaks before (T0) and after (T60) progestin exposure of T47D cells. **B)** Average enrichment and heatmap of PARP1 ChIP-seq peaks from panel A. Top is the average PARP1 enrichment for clusters T0, common and T60. Bottom heatmap for same three clusters spanning a 4kb regions of peaks center. Color scale represents the −log10-pval of PARP1 enrichment. **C)** Cistrome-dbtoolKit (Zheng et al., 2019) analysis for cluster T0 and common together and T60. The dbtoolKit results are shown as word clouds where the size of the name of the factor is proportional to the number of ChIP-seq experiments and the colors represent the different threshold numbers of retrieved experiments. **D)** Venn diagram showing the fraction of SEs found in cluster T0 + common and HCRs in cluster T60. **E**) Rank ordering of super-enhancers (ROSE) (Loven et al., 2013, Whyte et al., 2013) analysis for PARP1 ChIP-seq. PARP1 captures 471 SEs with key T47D genes associated with known SEs from the Super-Enhancer Data Base (SEdb) (Wang et al., 2023). **F)** Average enrichment and heatmap showing the distribution of PARP1 ChIP-seq for metagene analysis of T47D-SEs (the 471 calculated in panel E), SEs deposited in the SEdb and a control of SEs keratinocytes from SEdb. S=start, E=end of the genomic regions. Average enrichment represented as −log10-pval. **G)** Genome browser view of the representative GSE1 locus associated with a SE annotated in the SEdb (blue rectangles). Reported are the PARP1, BRD4, and H3K27ac ChIP-seq in T47D cells at both T0 and T60. ROSE predicted SE-PARP1 and SE-H3K27ac are indicated by the orange and green rectangles respectively. **H)** Average enrichment and heatmap of Progesterone Receptor (PGR) ChIP-seq across clusters from panel B. Color scale is indicated and represents the −log10-pval. **I)** Average enrichment and heatmap of PARP1 ChIP-seq for metagene analysis of 212 HCRs found in T47D (Le Dily et al., 2019b), and a random control of 212 regions across the genome. S=start, E=end of the genomic regions. Color scale is indicated and represents the −log10-pval. **J)** Genome browser view of the representative GREB1 locus associated with an HCR (indicated as purple rectangle). For this locus is reported also PGR ChIP-seq in T47D cells at both T0 and T60. For PGR T60 ChIP-seq two different progestin concentrations are shown (50pM and 10nM). Recruitment of PGR to the GREB1-HCR increases PARP1 occupancy as well as that of H3K27ac and BRD4.

PARP1 has also been proven to play a critical role in progesterone receptor (PGR) activity (Wright et al., 2012). Our clustering of PARP1 ChIP-seq (Figure 1B) and ChIP-seq enrichment analysis (Figure 1C), predicted cluster T60 to be bound by PGR. To prove this, we used available PGR ChIP-seq (Zaurin et al., 2021) at T0 and T60 and calculated its enrichment in our clusters (Figure 1H). As expected T60 cluster showed the greatest occupancy of PGR compared to the other two clusters (Figure 1H). If PGR is highly enriched at HCRs (Le Dily et al., 2019b) and PARP1 occupies 80% or HCRs, we investigated the chromatin binding dynamics of PARP1 at these regions before and after hormone exposure as we would expect increased PARP1 binding at T60. Indeed, we find by comparing PARP1 and BRD4 enrichment at HCRs that PARP1 showed a marked increase at T60 (Figure 1I) compared to the BRD4, which didn’t show significant changes (Figure 1I). PARP1 also showed a higher enrichment at HCRs than BRD4, which was considered a specific HCRs marker (Le Dily et al., 2019b). The *GREB1* gene is highly induced by progesterone (and progestin R5020) exposure and is associated with an HCR (Le Dily et al., 2019b) (Figure 1J). Hormonal-induced PGR occupancy at GREB1 locus was also linked to a surge in PARP1 occupancy at the GREB1-associated HCR (Figure 1J).

Albeit our analysis has focused on HCRs, our data shows that there are regions of PGR binding which are not HCR (non-HCRs) and that these are also targets of PARP1 occupancy upon hormone exposure. One example is the increased PARP1 binding at the intronic HAPRB within the *STAT5A* gene at T60 (Supp. Figure S1E) revealing a new regulatory region of *STAT5A* characterized by HCR-independent PARP1 binding. Therefore, all together our data show that PARP1 dynamically occupies chromatin in T47D breast cancer cells depending on signaling stimuli (such as progesterone) and can be considered as a new bona fide marker of T47D-SEs.

### PARP1 controls expression of SEs- and HCR-associated genes

If considered as a SEs marker, PARP1 should therefore be associated with highly expressed, cell type-specific genes normally controlled by SEs (Wang et al., 2019). To investigate whether PARP1 plays a crucial role in maintaining higher levels of transcription for SE-associated genes, we used RNAi to abrogate T47D PARP1 levels at T0 and T60 (Figure 2A and S2A) and used RNA-seq to probe the transcriptome in the absence of PARP1. In general, upon PARP1 KD at T0, a larger set of genes was down-regulated (1656) (compared to 1305 up-regulated) (Supp. Figure S2B) as expected based on the recently disclosed positive role of PARP1 in maintaining global nuclear chromatin accessibility (Ishii et al., 2024). When we plotted the absolute levels of gene expression per condition, we noticed that several highly expressed outlier genes did show a decrease in their transcript levels (Supp. Figure S2C). Therefore, we sought to determine whether PARP1 KD was indeed affecting predominantly genes with the highest absolute expression levels. To answer this, we first calculate absolute levels of all T47D-expressed genes at T0 and ranked them from high to low expression. We then divided all ranked genes into five classes and plotted the median expression for all five classes together with the median expression of PARP1-associated genes identified through Genomic Regions Enrichment of Annotations Tool (GREAT) (McLean et al., 2010) (Figure 2B). Our data shows that PARP1-associated genes possess higher absolute RNA levels, both at T0 and T60 (Figure 2B). We then selected the first 500 most highly expressed genes at T0 in our RNA-seq (with p-value < 0.05) and used hierarchical clustering to visualize their change in expression upon PARP1 KD at T0 and T60 (Figure 2C). Almost 70% of these 500 genes were significantly decreased in their expression (Figure 2C) both at T0 and T60 upon siPARP1. Among these genes we found several ones predicted to be associated with PARP1-SEs such as *KRT8, FOXA1, XBP1, KRT23, SCD, TOB1, TFRC, GSE1, PRLR and TRPS1* (Figure 2C) in accordance with our previous analysis (Figure 1E). We then asked whether, globally, genes associated with PARP1-SEs were downregulated and found that, collectively, they were significantly downregulated both at time T0 and T60 (Figure 2D). Three clear examples were the *TRSP1*, *PRLR and TOB1* whose transcript levels were profoundly affected by PARP1 KD prior and after progestin exposure (Figure 2E). Therefore, all together our data suggest that PARP1 is involved in the regulation of expression of SE-associated genes in T47D breast cancer cells.

**Figure 2.**
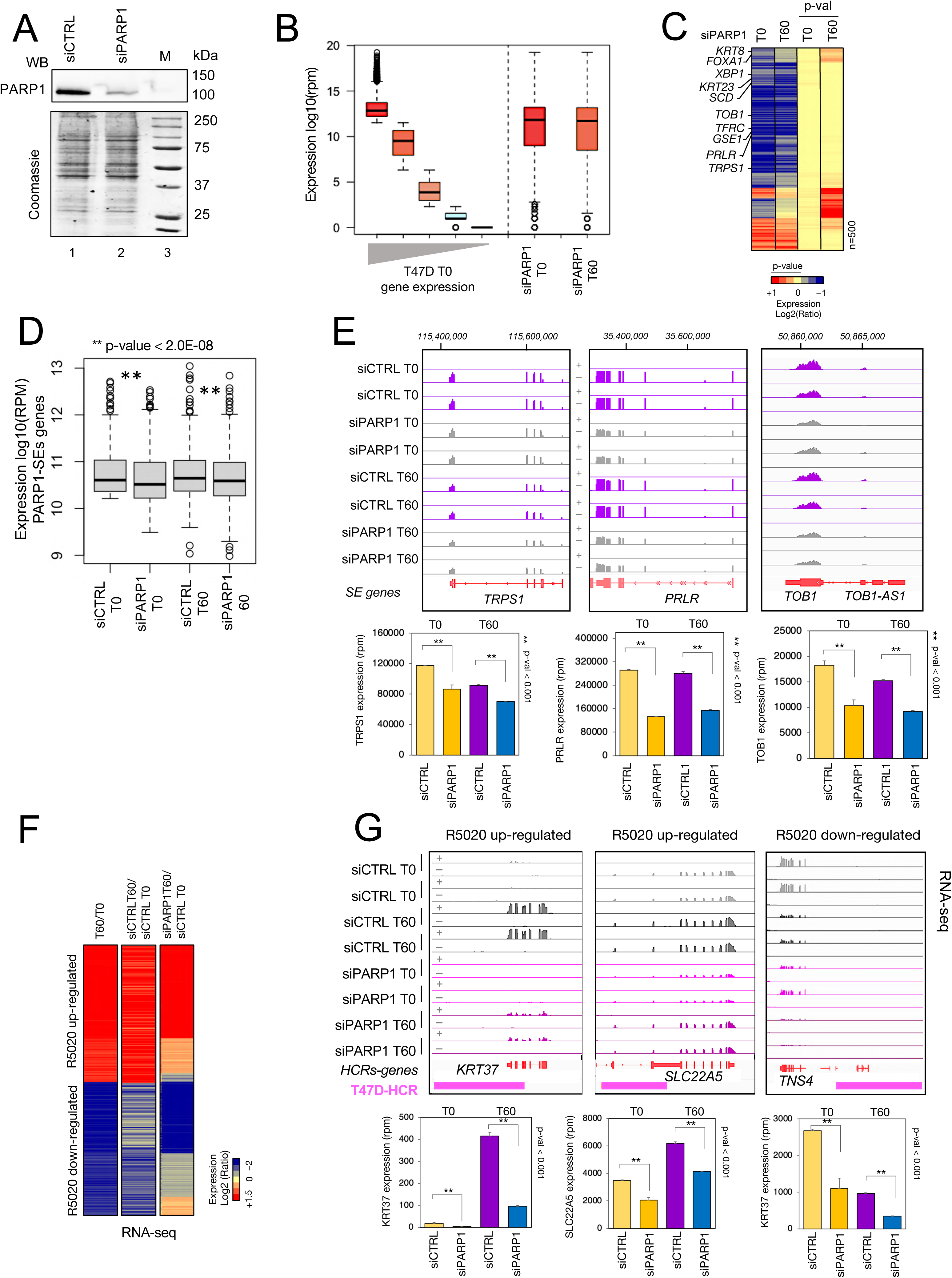
PARP1 knock down impairs gene expression of SE- and HCRs-associated genes. **A)** Western blot for PARP1 knock down (PARP1 KD) using siRNA. Indicated are the siRNA control scramble (siCTRL) and the siRNA against PARP1 (siPARP1). Marker with known molecular weight (250 to 25 kDa) proteins is also reported. The top panel displays the western blot for PARP1 protein. Bottom panels show Coomassie staining as loading control for the gel. **B)** Boxplot of median absolute expression levels of five groups of genes (all having the same number of genes) ranked based on their expression levels from RNA-sequencing of T47D at T0, from high (red) to low (cyan). The grey triangle highlights the progressive decrease in expression levels. Plotted is also the median absolute expression levels of genes associated with PARP1-SEs (siPARP1 T0 and siPARP1 T60 conditions). **C)** Heat map of first 500 genes (p-value < 0.05) ranked based on their absolute expression levels from the siPARP1 T0 RNA-seq. Reported are the changes in expression as log2 ratio of siPARP1 vs siCTRL at T0 and T60. For each gene p-values are also displayed for both T0 and T60 conditions. Among the significantly repressed genes are the PARP1-SEs associated genes such as *KRT8*, *FOXA1*, *XBP1*, *KRT23*, *SCD*, *TOB1*, *TFRC*, *GSE1*, *PRLR* and *TRPS1.* Color bar is indicated and represents the log2 ratio from +1 to −1 for gene expression and 0 to1 for p-values. **D)** Boxplot analysis of absolute expression of all PARP1-SEs associated genes in all conditions (siCTRL T0, siPARP1 T0, siCTRL T60 and siPARP1 T60). Two-tailed t-test p-values (**) are indicated for the pairwise comparisons of T0 and T60. **E)** Genome browser view (upper panel) and barplots (bottom panel) of RNA-seq tracks from data of two biological replicates of T47D cells treated with scramble siRNA (siCTRL) at both T0 and T60 (siCTRL T0, siCTRL T60) and siRNA against PARP1 (siPARP1) at both T0 and T60 (siPARP1 T0, siPARP1 T60). Shown are the representative TRPS1, PRLR and TOB1 loci associated with a PARP1-SE. **F)** Heatmap of R5020-(progestin) induced gene expression changes at T60 compared to R5020 expression changes upon PARP1 KD at T60. Color bar is indicated and represents the log2 ratio from +1.5 to −2. **G)** Genome browser view (upper panel) and barplots (bottom panel) of RNA-sequencing for the representative KRT37, SLC22A5 and TNS4 loci associated with an HCR (indicated as purple box).

As we find PARP1 also occupying HCRs, which control genes associated with hormonal response, we asked whether PARP1 abrogation might ultimately alter their expression. We therefore used published expression dataset (Le Dily et al., 2014) of R5020 exposure of T47D (T60) and calculated all the significantly differentially expressed genes compared to T0. We then generated an expression heatmap for both up- and down-regulated genes controlled by progestin in normal conditions and plotted the values for the same genes upon PARP1 ablation at T60 (Figure 2F). We also took scrambled siRNAs at time T60 (siCTRL T60) compared to the siCTRL at T0 (siCTRL T0) as a control for the use of siRNA, which could potentially interfere with R5020 action (Figure 2F). We find that samples in which PARP1 was abrogated resulted in loss of proper regulation of ∼45% of progestin-regulated genes compared to the normal condition (Figure 2F). siCTRL samples did not show significant alteration of the gene expression pattern induced by the hormone treatment (Figure 2F). Examples of hormone-regulated genes associated with HCRs such as *KRT37*, *SLC22A5* and *TNS4* clearly demonstrate the impact of PARP1 ablation on their response to progestin (Figure 2G). Altogether, our data show that PARP1 plays an important role in regulating genes associated with HCRs responding to progestin exposure.

### PARP1 maintains long-range chromatin loops for functional SEs and HCRs

To sustain high expression levels of cell type-specific genes, SEs use “looper” proteins to create genomic loops within the 3D structure of the human genome (Whyte et al., 2013). PARP1 has many features of a putative looper (Weintraub et al., 2017), being ubiquitously expressed within tissues, being a zinc-coordinating protein that binds hypo-methylated DNA sequences and capable of homodimerization (Weintraub et al., 2017). Moreover, PARP1 has been shown to act in stabilizing CTCF occupancy and maintaining open chromatin at active promoter during type III latency of Epstein-Barr Virus (EBV) (Lupey-Green et al., 2018). Therefore, it might be also important for looping of the human genome, favoring association of genes promoters to SEs and HCRs. Hence, we hypothesized that PARP1 could play a crucial role in establishment of chromatin looping involving SEs, HCRs and promoters, both as a direct “looper” or by assisting known architectural proteins. To address this hypothesis, we implemented Hi-C to first compare the 3D genome structure of T47D cells with WT or siRNA-silenced PARP1 First, we noticed that drastically reducing PARP1 levels did not alter the total number of topologically associating domains (TADs) (Supp. Figure S3A). Indeed, when comparing TAD numbers between siCTRL and siPARP1 Hi-C experiments (both at T0 and T60) we could not detect significant changes (Supp. Figure S3A). Also, when we calculated the insulation score across TAD boundaries, we found a slight decrease upon siPARP1 especially at T60 (Supp. Figure S3B), which could explain the largest decrease in TADs number across conditions (Supp. Figure S3A). A similar trend was observed for domains following PARP inhibitor Olaparib treatment which caused a slight decrease in the insulation score (Supp. Figure S3C). We then ensured that control siRNAs (siCTRL) did not alter the structure of the 3D genome, as shown for the TRPS1 locus (Supp. Figure S3D) and comparing insulation scores between siCTRL T0 and T0 (Supp. Figure S3B and S3C). However, when we measured long-range chromatin interactions or intra-TADs looping for TRPS1 (SE) and ZMYND8 (HCR) loci, PARP1 knock-down cells showed severely compromised looping structure, both at T0 (Figure 3A-B) as well as at T60 (Figure 3C-D). Albeit, the overall TAD structure is preserved upon PARP1 ablation, virtually all loops and stripes within the TAD disappeared (Figure 3A-C). This might explain the decrease in TRPS1 mRNA expression upon PARP1 depletion (Figure 2E), due to lack of TRPS1 chromatin looping necessary to maintain high levels of gene transcription. We then identified TAD boundaries in condition using identical algorithm settings (Ramirez et al., 2018, Wolff et al., 2018, Wolff et al., 2020), and we measured genomewide intra-TADs looping using Aggregate TAD Analyses (ATA) (van der Weide et al., 2021). At T0 siPARP1 already showed changes in intra-TADs looping (Figure 3E) but the strongest effect was obtained at T60 comparing the intra-TADs following progestin in the presence or absence of PARP1 (Figure 3F). Therefore, PARP1 KD at T0 caused a decrease in intra-TADs looping and a strong increase in the short close-range contact across the diagonal (Figure 3E, third panel), suggesting that the chromatin is becoming more compact (Ishii et al., 2024) with PARP1 having a role in regulating or limiting these local interactions. In contrast, at T60 we show that, apart from the decrease in contacts within TADs and at the corners, there was a strong decrease of the close-range interactions (Figure 3F, third panel). At T60 CTCF increased its chromatin-bound fraction upon siPARP1 (Supp. Figure S3E). We then inhibited PARP1 catalytic activity, both at T0 and at T60, and we observed an overall increase in intra-TADs looping, at the corners and over the diagonal (Supp. Figure SF-I). This suggests that the physical presence of PARP1 is important for long-range and short-range looping and that its PARylation might fine tune local interactions. To get a genomewide impression of looping we also performed the Aggregate Peak Analysis (APA) (van der Weide et al., 2021) on the set of loops we calculated for each condition (Wolff et al., 2022). Already at T0 siPARP1 shows decreased looping, which becomes more evident at T60 (Figure 3G-H), attesting to a more important genomewide role of PARP1 in looping upon hormone exposure. Reversely, Olaparib treatment showed higher levels of looping from T0 to T60 both with ATA (Supp. Figure S3F-G) and APA analysis (Supp. Figure S3H-I), arguing for a role of PARylation in restricting looping formation. As CTCF is the master regulator of chromatin looping, we therefore expected it to be highly enriched both at PARP1 peaks as well as at the promoters of genes down- and up-regulated by the PARP1 KD. However, when we plotted the enrichment of PARP1 clusters (from Figure 1B) for several factors (including PGR, H3K18ac, H3K27ac, FOXA1, PARP1, CTCF and POL2) CTCF only showed a very modest level of occupancy, compared to H3K27ac, PARP1 and POL2 (Figure 3I). Similar profiles were observed when we looked at genes down- and up-regulated by the PARP1 KD (siPARP1-down and siPARP1-up genes) (Figure 3J), where only H3K27ac, PARP1 and POL2 showed high enrichment within PARP1 regulated gene promoters, with again virtually no enrichment in CTCF (Figure 3J). Moreover, global levels of CTCF loading on chromatin were not affected by siPARP1at T0 (Figure 3K), suggesting that PARP1 could be one of the main proteins involved in looping between SE and siPARP1-down genes (which largely correspond to SE-genes) (Figure 2C, 3J). These findings motivated us to prove that mainly PARP1 (and not CTCF) was involved in maintaining the connection between the promoter of siPARP1-affected genes and the PARP1 peaks at the enhancers at T0. To answer this, we first calculated loops and separated the loop with and without CTCF anchor (Figure 3L). Comparing conditions, siPARP1 T0 and siPARP1 T60 loops were decreased compared to the siCTRL (Figure 3L), independently of a CTCF anchor, meaning that the looping structure is not unequivocally dictated by CTCF. Moreover, we generated aggregation plots at T0 comparing PARP1 peaks devoid of CTCF (ΔCTCF-PARP1), within enhancer regions, with siPARP1-down (Figure 3M, upper panels) and siPARP1-up genes (Supp. Figure S3J). CTCF peaks lacking overlap with PARP1 (CTCF-ΔPARP1) showed a weak looping and not a strong decrease upon siPARP1 T0 (Figure 3M, lower panel and Supp. S3J, lower panel), providing additional evidence to support a major role for PARP1 in connecting enhancers with PARP1 regulated genes. We also compared the PARP1 peaks overlapping with CTCF (CTCF-PARP1) which showed a strong decrease in looping upon PARP1 ablation (Supp. Figure S3K-L). If PARP1 is responsible for looping of enhancers and SEs to its own regulated genes we asked if this was the case also for HCRs. To answer this, we performed the same aggregation analysis at T60 by looking at PARP1 peaks overlapping HCRs, but devoid of CTCF. At T60 upon hormone exposure PARP1 relocates to PGR-bound regions (Figure 1B) and looping (Figure 3N, upper panel) was strong. Upon siPARP1 at T60 looping was dramatically decreased (Figure 3N, lower panel), suggesting that PARP1 participates in looping regions necessary for progesterone response (Figure 3N). All together these data suggest a role of PARP1 in linking SEs and HCRs to their respective genes via long-range looping (with PARylation playing instead a role in fine tuning short-range cis interactions) especially during progesterone exposure in T47D breast cancer cells. This is largely independent from CTCF which increased its chromatin-bound fraction upon siPARP1 at T60 suggesting a potential rescue mechanism to guarantee loading of looping factors after loss of chromatin-bound PARP1.

**Figure 3.**
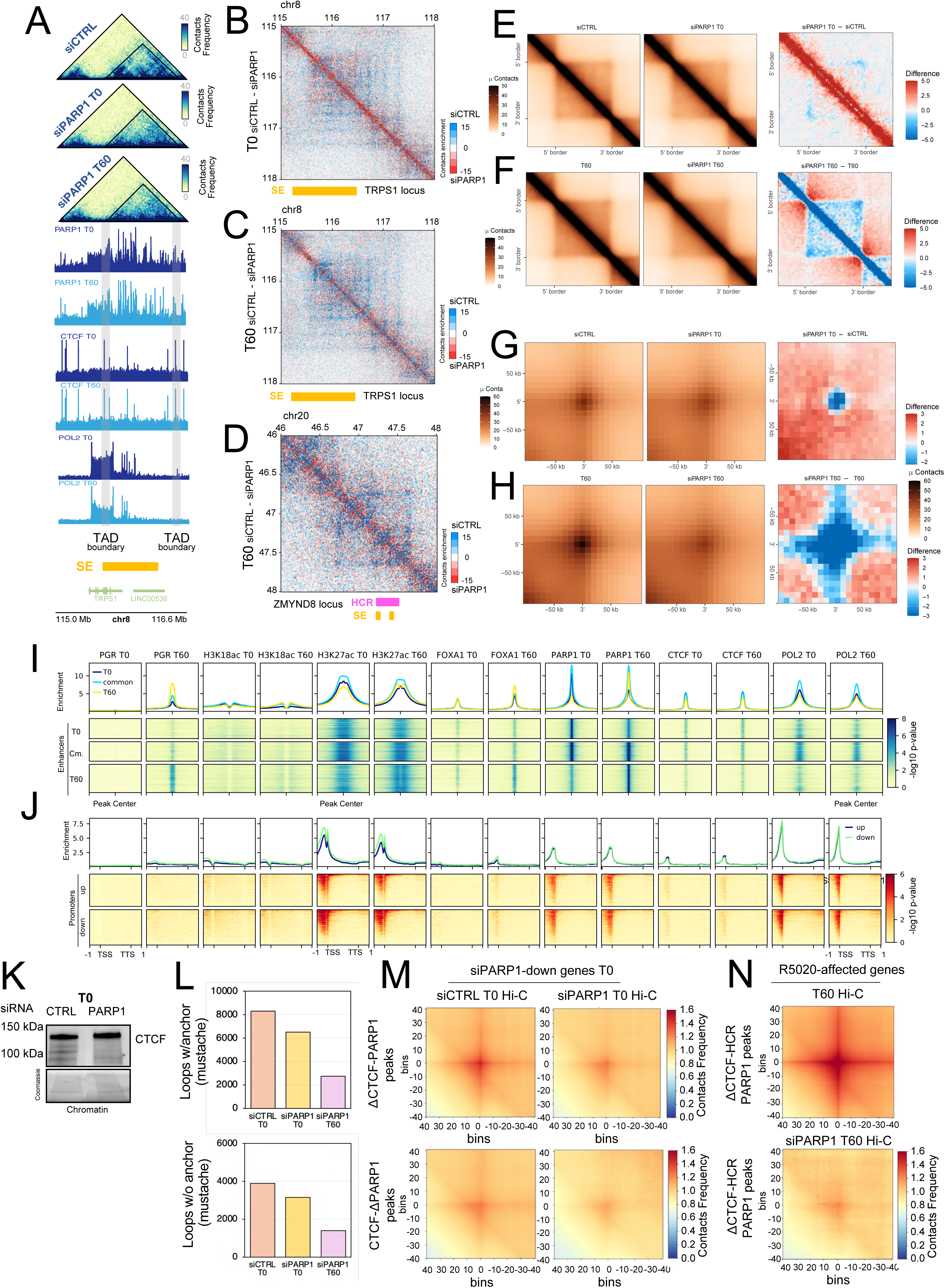
PARP1 is involved in long-range chromatin looping. **A)** Comparison of Hi-C matrix of contact frequency from siCTRL and siPARP1 for the TRPS1 locus. Represented is a region spanning almost 2Mb containing the *TRPS1* gene and its SE (indicated at the bottom in yellow). The Hi-C matrix is aligned with ChIP-seq of PARP1, CTCF and POL2 at T0 and T60 (middle portion of the panel). TADs are highlighted by the black triangle within the map and their boundaries by the grey rectangles. **B)**. Differential matrix plot for TRPS1 locus containing a SE at T0. **C)** Differential matrix plot for TRPS1 locus containing a SE at T60. For both differential plots TRSP1-SE is indicated by the orange rectangle. **D)** Differential matrix plot for ZMYDN8 locus containing an HCR (purple rectangle) at T60. For all matrix plots siPAR1 is subtracted from siCTRL. Indicated is the scale bar for contact enrichment towards siCTRL (blue) or siPARP1 (red). **E)** Normalized aggregate Hi-C contact heatmaps and differential contact heatmaps around rescaled T0 TADs (*n* = 4,072) in siCTRL and siPARP1 at T0. **F)** Normalized aggregate Hi-C contact heatmaps and differential contact heatmaps around rescaled T60 TADs (*n* = 4,167) in T60 and siPARP1 T60. Analysis and visualization of aggregate Hi-C contact heatmaps were performed using GENOVA’s ATA (van der Weide et al., 2021). **G**) Normalized aggregate peak analysis (APA) and differential aggregate peak heatmaps around T0 calculated loops (Wolff et al., 2022), comparing siCTRL and siPARP1 at T0. **H)** Normalized aggregate peak analysis (APA) and differential aggregate peak heatmaps around calculated T60 loops (*n* = 4,231) at T60 and siPARP1 T60. Normalized aggregate peak analysis has been performed using GENOVA’s APA (van der Weide et al., 2021). **I)** Heatmap showing enrichment for PGR, H3K18ac, H3K27ac, FOXA1, PARP1, CTCF and POL2 (at both T0 and T60) over clusters from figure 1B devoid of promoters (only distal cis-regulatory peaks were considered “Enhancers”). H3K27ac, PARP1 and POL2 show high enrichment. **J)** Heatmap showing enrichment for same factor indicated in panel I over genes down- and up-regulated upon PARP1 KD at T0. H3K27ac, PARP1 and POL2 show high enrichment. **K)** Western blot assessing chromatin loading of CTCF in T47D treated with siRNA CTRL and PARP1 at T0. No difference in levels of CTCF can be seen on chromatin. **L)** Chromatin loops across T0, siCTRL T0 and siPARP1 T0 conditions calculated using mustache (Roayaei Ardakany et al., 2020). Upper panels are total loops, bottom panel loops without CTCF anchors. **M)** Aggregate plots (Ramirez et al., 2018, Wolff et al., 2018, Wolff et al., 2020) of Hi-C interactions between PARP1 peaks devoid of CTCF (ΔCTCF-PARP1, upper figure) and CTCF devoid of PARP1 (ΔCTCF-PARP1, lower figure) with genes down-regulated by PARP1 KD (siPARP1-down genes T0). Compared are the Hi-C data of siCTRL and siPARP1 at T0 for both sets of peaks. ΔCTCF-PARP1 peaks showed higher levels of interaction with siPARP1-down genes T0. **N)** Aggregate plots of Hi-C interactions between PARP1 peaks at HCRs devoid of CTCF (ΔCTCF-HCR PARP1) and genes affected by R5020 (from Figure 2G). Compared are the Hi-C data of T60 and siPARP1 at T60. Note the strong enrichment at T60 for these regions as previously reported (Le Dily et al., 2019b).

### PARP1 ablation but not its catalytic activity affects chromatin compartments

Given the strong correlation between PARP1 and H3K27ac we showed here (Supp. Figure S1B) and the known roles of PARP1 in regulating heterochromatin (Dantzer and Santoro, 2013, Guetg and Santoro, 2012) and orchestrating epigenetic events setting up chromatin domains (Ciccarone et al., 2017), we speculated that PARP1 could also be involved in regulating chromatin compartments. To investigate this, we calculated A and B compartments for siCTRL and siPARP1 at T0 and use saddle plots to show global changes in chromatin compartments (Figure 4A). siPARP1 at T0 showed a marked decrease in A-A and B-B compartment association (Figure 4A) and loss of compartment strength for almost all chromosome arms (Figure 4B). It also seems that PARP1 KD had a most pronounced effect on the disappearance of A compartments (Figure 4A, differential plot, right panel). Indeed, when looking at chr8 and selecting regions with changes in compartments (highlighted), we could only detect loss of various A compartments (Figure 4C, shaded in grey), such as the one comprising the TRPS1 locus. This was also true for the PRLR locus which showed A to B conversion upon siPARP1 at both T0 and T60 (Figure 4D). PARP1 KD showed whole chromatin compartment changes at the chromosome levels (Figure 4E). To test the role of PARylation, we calculated A and B compartments for Olaparib treatment of T47D cells at T0 and use saddle plots to show global changes in chromatin compartments (Figure 4F). Olaparib at T0 did not show marked changes in in A-A and B-B compartment association (Figure 4F) and virtually no changes in compartment strength for almost all chromosome arms (Figure 4G). We find that even at T60 Olaparib treatment did not significantly affect either compartment interactions (Figure 4H) nor compartment strength (Figure 4I). All together these data shows that PARP1, but not its catalytic activity, plays a role in regulation of chromatin compartmentalization.

**Figure 4.**
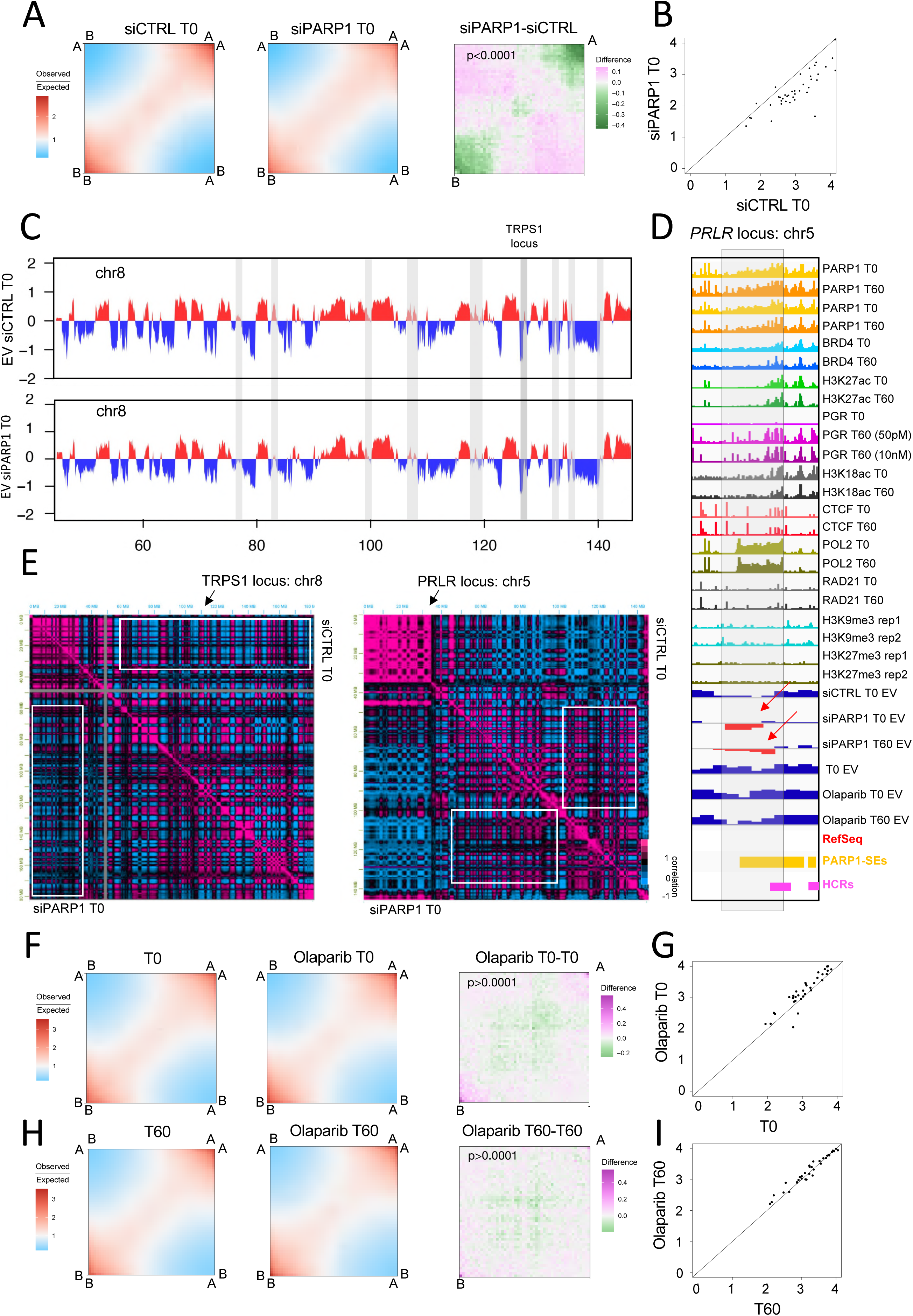
Compartments interactions depend on PARP1 and its catalytic activity. **A)** GENOVA (van der Weide et al., 2021) saddle plot analysis of siCTRL and PARP1 compartments interaction. The difference between the two samples. Upon siPARP1 there is a decrease interaction between compartments, especially A-A. **B)** GENOVA compartments strength analysis for chromosome arms. The per-arm compartment strength is at almost every arm lower in the PARP1 KD. **C)** GENOVA comparison of Eigen Vectors for chromosome 8 (containing the TRPS1-SE locus as indicated) for siCTRL and siPARP1 Hi-C samples. Grey rectangles indicate the loci of change in compartments. Predominantly we observe a reduction in A compartments or a A to B switch, as at the TRPS1 locus. **D)** Genome browser view of the PRLR locus with all the indicated ChIP-seq of histone marks and proteins. Reported are also the eigen vectors (EV) calculated for all the conditions tested: siCTRL T0, siPARP1 T0, siPARP1 T60, T0, Olaparib T0 and Olaparib T60. Only the KD of PARP1 induces at this locus an inversion of compartment from A to B (indicated by the red arrows). **F)** GENOVA (van der Weide et al., 2021) saddle plot analysis of T0 and T0 Olaparib compartments interaction. Showed is also the difference between the two samples. Upon Olaparib treatment at T0 there is a slight increase interaction between compartments, especially B-B. **G)** GENOVA compartments strength analysis for chromosome arms. The per-arm compartment strength is not really affected by T0 Olaparib treatment. **H)** Saddle plot analysis of T60 and T60 Olaparib compartments interaction. The difference between the two samples is also shown. Upon Olaparib treatment at T60 there is no changes in interaction between compartments. **I)** GENOVA compartments strength analysis for chromosome arms. The per-arm compartment strength is not really affected by T60 Olaparib treatment.

## Discussion

The multifaceted roles of PARP1 in transcriptional regulation in breast cancer cells have been further elucidated through our current findings. Here we expand the known functions of PARP1 beyond its well-established involvement in DNA repair (Ray Chaudhuri and Nussenzweig, 2017), highlighting its role at Super-Enhancers (SEs), and its dynamic redistribution to Hormone-Control Regions (HCRs) in response to progestin exposure and its involvement in intra-TAD and chromatin compartment organization.

Our ChIP-seq of PARP1 in T47D cells shows a dynamic genomewide occupancy with a switch of PARP1 binding, following exposure to progestin, towards regions bound by PGR (comprising most HCRs). Our data also show that occupancy of PARP1 at SEs and HCRs is directly correlated with their associated genes transcription. Indeed, PARP1 ablation by siRNA caused major changes in the gene expression of these genes. Prior studies have established that SEs are critical for the transcriptional control of genes involved in cell identity and disease progression, including cancer (Whyte et al., 2013). The dynamic redistribution of PARP1 from SEs to HCRs upon progestin exposure unequivocally proves a direct involvement of PARP1 in mediating hormonal responses at the chromatin level. This observation aligns with the growing body of evidence indicating that PARP1 is not only a DNA damage sensor but also a key regulator of transcriptional programs (Krastev et al., 2018). Our data further support this notion by showing that PARP1 controls the expression of both SE- and HCR-associated genes, thus influencing cellular phenotypes linked to hormone responsiveness.

Moreover, we provide evidence that the control exerted by PARP1 on SEs and HCRs is mediated by the creation of long- and short-range chromatin loops capable of connecting SEs and HCRs with their respective genes. When PARP1 is knocked down before (T0) and after (T60) progestin exposure, there is a decrease in genome-wide looping, particularly more pronounced at T60. Hence our results indicate that PARP1 plays an essential role in maintaining long-range chromatin loops necessary for the functionality of SEs and HCRs. This finding resonates with recent advances in 3D genome organization, which have revealed that transcriptional enhancers often communicate with target promoters via chromatin looping (Dekker and Mirny, 2016). The disruption of these loops upon PARP1 ablation underscores the importance of PARP1 in orchestrating higher-order chromatin architecture, a process that appears to be independent of its catalytic activity. This distinction is particularly intriguing given the widespread use of PARP1 inhibitors as therapeutic agents in cancer treatment (Lord and Ashworth, 2017). Our results suggest that the non-catalytic functions of PARP1 may also be relevant for its oncogenic roles, potentially opening new avenues for therapeutic intervention.

As several evidence are accumulating regarding the complex universe of inactive PARP1 (Huang et al., 2024, Matveeva et al., 2022), our data suggest that PARP1 might play a structural role in maintaining or promoting chromatin looping interactions across the genome independently of its catalytic activity. The decrease in looping upon PARP1 knockdown indicates that PARP1’s presence is necessary for the formation or stabilization of these loops, especially as the cells respond to progesterone over time.

The stronger effect at T60 might imply that PARP1’s role in chromatin looping becomes more critical as cells adapt to progesterone exposure which reflects a dynamic role where PARP1’s influence on genome structure evolves with hormonal exposure and its dynamic change in genomewide occupancy. When PARylation (the catalytic activity of PARP1) is inhibited, there’s an increase in genome-wide looping, mirroring the effects seen with WAPL knockout (Haarhuis et al., 2017). WAPL is known to be involved in the removal of cohesin from chromatin, which leads to the dissolution of loops. By inhibiting PARylation, you remove this regulatory mechanism, leading to an increase in looping, suggesting that PARylation by PARP1 might normally act to limit or fine-tune chromatin interactions, possibly creating a less favorable environment for loop formation by modifying proteins that interact with chromatin or by altering the chromatin itself.

The study by Lupey-Green et al. shows that PARP1 inhibition decreases CTCF binding to the EBV genome, which contrasts with the increase in looping observed in our experiments. CTCF is a key protein in forming and stabilizing chromatin loops, often acting as an anchor for cohesin (van Ruiten and Rowland, 2021). The decrease in CTCF binding might suggest that PARP1’s role in loop regulation could be context-dependent having different roles in different genomic environments or cell types and it may be dependent on PARP1’s own post-translational modifications. In the context of EBV, PARP1 might support CTCF binding, which in turn supports loop formation. However, in the broader genomic context, especially in response to progesterone, PARP1’s enzymatic activity might restrict looping through mechanisms not involving CTCF directly. While PARylation might generally limit loop formation by modifying proteins or chromatin, in the case of EBV, this modification might be crucial for CTCF’s role, suggesting a nuanced interaction where PARP1’s activity might enhance or diminish loop formation based on the specific protein interactions.

Ultimately, given the very well-known role of PARP1 in DNA repair, it might not be too surprising locating this protein to SEs and HCRs as genome-wide mapping of DSBs revealed an unforeseen coupling mechanism between transcription and DNA repair at super-enhancers, as means of hyper-transcription of oncogenic drivers (Oster and Aqeilan, 2020, Hazan et al., 2019).

In summary, our study provides novel insights into the multifaceted role of PARP1 in breast cancer cells, emphasizing its involvement in Super-enhancer- and hormone-mediated transcriptional regulation together with chromatin organization. These findings expand on existing knowledge by revealing a dynamic interplay between PARP1 and distinct genomic elements, such as SEs and HCRs, and highlight the potential for targeting PARP1 beyond its canonical catalytic functions. Future investigations should aim to elucidate the molecular mechanisms underlying PARP1’s non-catalytic activities and explore their implications for breast cancer therapy.

## STAR METHODS

### Key resources table

| REAGENT or RESOURCE | SOURCE | IDENTIFIER |
| --- | --- | --- |
| **Antibodies** |  |  |
| Rabbit polyclonal antibody anti-CTCF | Active Motif | Cat#61311; RRID: AB_2614975 |
| Rabbit polyclonal antibody anti-PARP-1 N-terminal | Active Motif | Cat#39559; RRID: AB_2793257 |
| StarBright Blue 700 anti-rabbit | Biorad | Cat#12004161 |
| **Chemicals, peptides, and recombinant proteins** |  |  |
| RPMI1640 RED phenol | Thermo Fisher Scientific | Cat#11875093 |
| RPMI 1640 No Phenol Red | Thermo Fisher Scientific | Cat#11835030 |
| Fetal Bovine Serum | Gibco | Cat#A5256701 |
| Fetal Bovin Serum, charcoal stripped | Thermo Fisher Scientific | Cat#12676029 |
| 0,05% Trypsin-EDTA 1x | Gibco | Cat#25300-054 |
| L-Glutamine 200 mM 100x | VWR | Cat#392-0441 |
| Penicillin-Streptomycin solution 100x | EuroClone | Cat#ECB-3001D |
| Lipofectamine 3000 | Invitrogen | Cat#L3000001 |
| Lipofectamine 2000 | Thermo Fisher Scientific | Cat#11668019 |
| Dynabeads Protein A | Thermo Fisher Scientific | Cat#10008D |
| Dynabeads Protein G | Thermo Fisher Scientific | Cat#10003D |
| Halt Protease Inhibitor Cocktail 100X | Thermo Scientific | Cat#78438 |
| Complete EDTA free Protease Inhibitor Cocktail | Roche | Cat#11836170001 |
| Glycine | Merck | Cat#GE17-1323-01 |
| NP40/IGEPAL | Sigma-Aldrich | Cat#18896<br>CAS: 9002-93.1 |
| Triton-X-100 | Sigma-Aldrich | Cat#T8787 |
| Gel red | Sigma-Aldrich | Cat#SCT123 |
| RNAse A | Thermo Fisher Scientific | Cat#R1253 |
| Agarose | Sigma-Aldrich | Cat#A9539 |
| R5020 | MedChemExpress | Cat#HY-119384<br>CAS No: 34184-77-5 |
| Olaparib | Sigma-Aldrich | Cat#SML3705 |
| Formaldehyde | Thermo Fisher Scientific | Cat# 033314.AP<br>CAS: 50-00-0 |
| Skim Milk Powder | Sigma | Cat#70166-500G |
| Critical commercial assays |  |  |
| RNeasy Plus Mini Kit | Qiagen | Cat#74134 |
| ChIP-IT High sensitivity ChIP kit | Active Motif | Cat#53040 |
| Mini-PROTEAN TGX Stain-Free Protein Gels 4-20% | Bio-Rad | Cat#4568093 |
| Trans-Blot SD Cell | Bio-Rad | Cat#221BR-30822 |
| Protein Assay Dye Reagent | Bio-Rad | Cat#5000006 |
| **Deposited data** |  |  |
| RNA-seq siControl 2 replicates T47D T0 | This study | GSE291980 |
| RNA-seq siControl 2 replicates T47D T60 | This study | GSE291980 |
| RNA-seq siPARP1 2 replicates T47D T0 | This study | GSE291980 |
| RNA-seq siPARP1 2 replicates T47D T60 | This study | GSE291980 |
| PARP1 ChIP sequencing R5020 T0 T47D | This study | GSE291980 |
| PARP1 ChIP sequencing R5020 T60 T47D | This study | GSE291980 |
| siCTRL Hi-C sequencing T0 | (Ferrari et al., 2020) | GSE120162 |
| siPARP1 Hi-C sequencing T0 | This study | GSE291980 |
| siPARP1 Hi-C sequencing T60 | This study | GSE291980 |
| Olaparib-treated Hi-C sequencing T0 | This study | GSE291980 |
| Olaparib-treated Hi-C sequencing T60 | This study | GSE291980 |
| T0 Hi-C sequencing | (Ferrari et al., 2020) | GSE120162 |
| T60 Hi-C sequencing | (Le Dily et al., 2014) | GSE53463 |
| RNA Pol2 ChIP-sequencing T0 | (Ferrari et al., 2020) | GSE120162 |
| RNA Pol2 ChIP-sequencing T60 | (Le Dily et al., 2019b) | GSE109229 |
| H3K27ac ChIP sequencing R5020 T0 T47D | (Ferrari et al., 2020) | GSE120162 |
| H3K27ac ChIP sequencing R5020 T60 T47D | (Ferrari et al., 2020) | GSE120162 |
| H3K18ac ChIP sequencing R5020 T0 T47D | (Ferrari et al., 2020) | GSE120162 |
| H3K18ac ChIP sequencing R5020 T60 T47D | (Ferrari et al., 2020) | GSE120162 |
| BRD4 ChIP sequencing R5020 T0 T47D | (Le Dily et al., 2019b) | GSE109229 |
| BRD4 ChIP sequencing R5020 T60 T47D | (Le Dily et al., 2019b) | GSE109229 |
| PGR ChIP sequencing R5020 T0 T47D | (Zaurin et al., 2021) | GSE179666 |
| PGR ChIP sequencing R5020 T60 T47D 50pM | (Zaurin et al., 2021) | GSE179666 |
| PGR ChIP sequencing R5020 T60 T47D 10nM | (Zaurin et al., 2021) | GSE179666 |
| CTCF ChIP sequencing R5020 T0 T47D | (Le Dily et al., 2014) | GSE53463 |
| CTCF ChIP sequencing R5020 T60 T47D | (Le Dily et al., 2014) | GSE53463 |
| H3K9me3 ChIP-sequencing normal conditions T47D | (Ramírez-Cuéllar et al., 2024) | GSE247777 |
| H3K27me3 ChIP-sequencing normal conditions T47D | (Ramírez-Cuéllar et al., 2024) | GSE247777 |
| H3K36me3 ChIP-sequencing normal conditions T47D | (Zhang et al., 2016) | GSM1541451 |
| **Experimental models: Cell lines** |  |  |
| T47D breast cancer cell lines | ATCC | CRL-2865 |
| **siRNA/Oligonucleotides** |  |  |
| siGENOME ON-TARGET plus SMART pool against human *PARP1* siRNA | Dharmacon | L-00-6656-03-0010 |
| siGENOME Non-Targeting siRNA pool #2 | Dharmacon | D-001206-14-20 |
| **Software and algorithms** |  |  |
| Bowtie2 | (Langmead et al., 2009) |  |
| STAR | (Dobin and Gingeras, 2016) |  |
| Samtools | (Li et al., 2009) |  |
| Cistrome | (Zheng et al., 2019) |  |
| GENOVA | (van der Weide et al., 2021) |  |
| Plotgardener | (Kramer et al., 2022) |  |
| Deeptools | (Ramirez et al., 2016) | https://deeptools.readthedocs.io/en/latest |
| hicexplorer | (Wolff et al., 2020, Wolff et al., 2018) |  |
| HiC-Pro | (Servant et al., 2015) |  |
| ROSE | (Whyte et al., 2013, Hnisz et al., 2013) |  |
| Juicer | (Durand et al., 2016) |  |
| IGV | (Thorvaldsdottir et al., 2013) |  |

## Methods Details

### Cell Lines and Treatments

Human T47D cells (American Type Culture Collection [ATCC]: CRL-2865) were grown in RPMI supplemented with 10% fetal bovine serum (FBS); Cells were than treated prepared for hormone exposure culturing them in RPMI supplemented with 10% charcoal-treated FBS for 48 h and starvation was achieved by withdraw of FBS for 16h (referred as T0 condition in the figures and text). T0 T47D cells were subjected to R5020 (progestin) exposure at 10nM for 1h (referred as T60 condition in the figures and text).

### Antibodies

Rabbit polyclonal antibodies against CTCF (61311) and PARP1 (39559) were obtained from Active Motif and used for Western blotting. The secondary antibody, Goat anti-Rabbit IgG StarBright Blue 700 (12004161), was from Bio-Rad.

### PARP1 ChIP

PARP1 ChIP was carried out using ChIP-grade antibody AB_2793715 from Active Motif and High-sensitivity ChIP protocol (ChIP-IT High Sensitivity®, Active Motif). 25 µg (measure by qubit) of ChIP-IT High Sensitivity®-prepared chromatin were used in ChIP with 8 µl of PARP1 AB_27937 N-terminal antibody. DNA was purified using phenol/chloroform and precipitation and quantified using qubit dsDNA HS. 1 ng of DNA was used for library preparation. PARP1 ChIP was carried out in T47D for both T0 and T60 time points in duplicate (see GEO submission).

### ChIP-seq library preparation

Libraries were prepared using the NEBNext® Ultra DNA Library Prep for Illumina® kit (ref. E7370) according to the manufacturer’s protocol. Briefly, input and ChIP enriched DNA were subjected to end repair and addition of “A” bases to 3′ ends, ligation of adapters and USER excision. All purification steps were performed using AgenCourt AMPure XP beads (ref. A63882, Beckman Coulter). Library amplification was performed by PCR using NEBNext® Multiplex Oligos for Illumina (Index Primers Set 1, ref. E7335), (Index Primers Set 2, ref. E7500), (Index Primers Set 3, ref. E7710) and (Index Primers Set 4, ref. E7730). Final libraries were analyzed using Agilent Bioanalyzer or Fragment analyzer High Sensitivity assay (ref. 5067-4626 or ref. DNF-474) to estimate the quantity and check size distribution and were then quantified by qPCR using the KAPA Library Quantification Kit (ref. KK4835, KapaBiosystems) prior to amplification with Illumina’s cBot. Libraries were sequenced 1 * 50^+8^ bp on Illumina’s HiSeq2500.

### ChIP-seq analysis and Peak Calling

Analysis of sequence data was carried out as previously described (Ferrari et al., 2014) with minor modifications. Reads were aligned to the hg38 human genome reference sequence (GRCh38) using Bowtie (Langmead et al., 2009) and aligning parameters of uniqueness (-S –m1 –v2 –t –q). p-values for the significance of ChIP-seq counts compared to input DNA were calculated as described (Pellegrini and Ferrari, 2012) using a p-value threshold of 10^−8^ and a false discovery rate (FDR) < 1%.

### ChIP-seq Downstream Analysis

Average ChIP-seq signals of 50 bp windows around 1, 2 or 3 kb upstream and downstream of annotated peaks or TSSs were calculated using the deeptools (https://deeptools.readthedocs.io/en/latest/) (Ramirez et al., 2016), both using the computeMatrix function (reference-point for peaks centred analysis, or scale-regions for metaregion analysis).

### Deeptools

computeMatrix and plotHeatmap functions (Ramirez et al., 2016) have been implemented for generating all metagene averages and heatmaps reported in this paper.

### Cistrome Database and Cistrome-tookit

Cistrome toolkit analysis was performed using the “What factors have a significant binding overlap with your peak set” and selecting all peaks for transcription factors and chromatin regulators (Zheng et al., 2019). Resulted table in tsv have been analyzed for transcription factors and chromatin regulators by counting the number of times the name appeared and its calculated wiggle score. Based on these two parameters we implemented wordcound representation of the results as illustrated in figure 1C.

### RNA extraction and library preparation

Libraries were prepared using the NEBNext® Ultra DNA Library Prep for Illumina® kit (ref. E7370) according to the manufacturer’s protocol. Briefly, input and ChIP enriched DNA were subjected to end repair and addition of “A” bases to 3′ ends, ligation of adapters and USER excision. All purification steps were performed using AgenCourt AMPure XP beads (ref. A63882, Beckman Coulter). Library amplification was performed by PCR using NEBNext® Multiplex Oligos for Illumina (Index Primers Set 1, ref. E7335), (Index Primers Set 2, ref. E7500), (Index Primers Set 3, ref. E7710) and (Index Primers Set 4, ref. E7730). Final libraries were analyzed using Agilent Bioanalyzer or Fragment analyzer High Sensitivity assay (ref. 5067-4626 or ref. DNF-474) to estimate the quantity and check size distribution and were then quantified by qPCR using the KAPA Library Quantification Kit (ref. KK4835, KapaBiosystems) prior to amplification with Illumina’s cBot. Libraries were sequenced 1 * 50+8 bp on Illumina’s HiSeq2500.

### RNA-seq Pipeline and Differential Gene Expression Analysis

Sequencing adapters and low-quality ends were trimmed from the reads using Trimmomatic, using the parameters values recommended (Bolger et al., 2014) and elsewhere (https://goo.gl/VzoqQq) (trimmomatic PE raw_fastq trimmed_fastq ILLUMINACLIP:TruSeq3-PE.fa:2:30:12:1:true LEADING:3 TRAILING:3 MAXINFO:50:0.999 MINLEN:36). The trimmed reads were aligned to GRCh38 (Lander et al., 2001) using STAR (Dobin et al., 2013).

First, the genome index files for STAR were generated with: star–runMode genomeGenerate–genomeDir GENOME_DIR–genomeFastaFiles genome_fasta– runThreadN slots–sjdbOverhang read_length–sjdbGTFfile sjdb–outFileNamePrefix GENOME_DIR/

Where genome_fasta is the FASTA file containing the GRCh38 sequence downloaded from the University of California Santa Cruz (UCSC) Genome Browser, excluding the random scaffolds and the alternative haplotypes; and sjdb is the GTF file with the GENCODE’s V24 annotation.

Second, trimmed reads were aligned to the indexed genome with: star–genomeDir GENOME_DIR/–genomeLoad NoSharedMemory–runThreadN slots–outFilterType “BySJout”–outFilterMultimapNmax 20–alignSJoverhangMin 8–alignSJDBoverhangMin 1–outFilterMismatchNmax 999–outFilterMismatchNoverLmax 0.04–alignIntronMin 20– alignIntronMax 1000000–alignMatesGapMax 1000000–readFilesIn read1 read2– outSAMtype BAM SortedByCoordinate–outTmpDir TMP_DIR/–outFileNamePrefix ODIR1/$sample_id.–outWigType bedGraph–readFilesCommand zcat (https://docs.google.com/document/d/1yRZevDdjxkEmda9WF5-qoaRjIROZmicndPl3xetFftY/edit?usp=sharing)

Differences in gene expression were calculated by using a DESeq2. Genes with fold change (FC) ± 1.5 (p value < 0.05; FDR < 0.01) were considered as significantly regulated in the volcano plots. However, for highly expressed genes (such those associated with SEs), we considered only the p-value < 0.00001 if the lof2FC was negative, as small changes for these highly expressed genes could be significant even though the FC is not that high or small.

### Bedtools

Bed intersection was carried out using bedtools (Quinlan and Hall, 2010) “intersectBed” function with default parameters of 1-bp overlap. Graphic representation of Venn diagrams has been obtained with R graphic, using R-studio (https://www.rstudio.com).

### siRNA knockdown of PARP1

Dharmacon L-006656-03-0010 ON-TARGET plus SMART pool against human *PARP1* siRNA and D-001206-14-20 siGENOME non-targeting siRNA Pool #2 were used to carry out PARP1 knockdown in T47D cells. Cells were seeded in RPMI medium without phenol red supplemented with 10% dextran-coated charcoal-treated FBS (DCC*/*FBS) and culture for 16 h prior to transfection with lipofectamine (Lipofectamine 3000, Invitrogen). siRNAs were used at 40 nM and cells were left in culture for 48 h in the presence of the siRNA. Cells were subjected to serum starvation for 16 h prior to further processing. The knockdown efficiency was evaluated by SDS-PAGE/Western Blot analysis.

### Cellular Biochemical Fractionation

Protein fractions from cytoplasm, nucleoplasm, and chromatin were obtained as described (REF). A total of 1.2×10^6^ cells were washed on plate with cold PBS and harvest in 1ml PBS using a cell scraper; cells were spun down at 1000 rpm for 2 min and the supernatant discarded. The remaining cell pellet was washed twice with PBS. (For each wash cells were resuspended in 1ml cold PBS added of Halt Proteinase Inhibitor Cocktail (Thermo Scientific) and spun down at 1000 rpm for 2 min). Cell pellet was then resuspended in 100 μl of Buffer A (10 mM HEPES pH 7.9, 10 mM KCl, 1.5 mM MgCl2, 0.34 M Sucrose, 10% Glycerol, 1 mM DTT and Halt Proteinase Inhibitor Cocktail). Triton X-100 was then added to the cell pellet to a final concentration of 0.1% and incubated on ice for 8 min. After incubation, the resuspended cell pellet was centrifuged at 1,300 x g at 4°C, for 5 min; The resulting supernatant (named fraction S1) was separate from the pellet (corresponded to nuclei and named P1). S1 fraction was then clarified by high-speed centrifugation at 20,000 x g at 4°C, for 5 min; the resulting supernatant was collected and named fraction S2 (soluble cytoplasmic extract). The pellet (named P2) was discarded. P1 fraction was then washed once with 100 μl of Buffer A, spun down at 1000 rpm at 4°C for 2 min and lysed for 30 min in 30 μl of Buffer B (3 mM EDTA, 0.2 mM EGTA, 1 mM DTT and Halt Proteinase Inhibitor Cocktail). The resulting lysate was then centrifuged at 1,700 x g at 4°C, for 5 min. The supernatant corresponding to the soluble nuclear extract was collected (and named fraction S3) and separated from the pellet (insoluble chromatin) named fraction P3. P3 was washed one more time with 30 μl of Buffer B, centrifuged at 1,700 x g at 4°C, for 5 min and the resulting pellet (after discarding the supernatant) was resuspend in 50 μl SDS sample buffer (62.5 mM Tris-HCl pH 6.8, 2.5% SDS, 0.002% Bromophenol Blue, 0.7135 M β-mercaptoethanol and 10% glycerol) and boiled for 10 min at 70°C.

### Protein quantification and western blot

Protein Assay Dye Reagent (Biorad) was used to quantify fractions S1 and S3. 10 μg of each S3 fractions and 5μl of the P3 fraction were loaded for SDS-PAGE/Western Blot analysis. Pre-casted gels (4%–20% polyacrylamide) were used for all SDS-PAGE/Western analysis at room temperature (150V for 70 min). Proteins transfer to Immuno-blot PVDF membrane 0,2 μm was carried out using Towbin transferring buffer and TRANS-BLOT SD semi-dry transfer cell at RT (15V for 35 min). Membrane blocking was carried out using Skim Milk Powder (5% in TBS-T) at room temperature for 1 h. For western blots primary antibodies were used at 1:20000 for PARP1 (Active Motif, 39559) and 1:1000 for CTCF (Active Motif, 61311), incubated overnight at 4°C followed by 1 h incubation with StarBright Blue 700 anti-rabbit (12004161, Biorad) and blots were developed using ChemiDoc MP Imaging System (Biorad) according to the manufacturer instructions.

### In situ Hi-C library preparation

In situ Hi-C experiments were performed as previously described (Rao et al., 2014) with some modifications (Le Dily et al., 2019a). Adherent cells were directly cross-linked on the plates with 1% formaldehyde for 10 min at room temperature. Reaction was stopped with glycine (125 mM final) and cells were recovered by scrapping in PBS. Cross-linked cells were lysed 30 min on ice in 3C lysis buffer (10 mM Tris-HCl at pH 8, 10 mM NaCl, 0.2% NP-40, 1× anti-protease cocktail) and resuspended in 190 µL of NEBuffer2 1× (New England BioLabs [NEB]) after centrifugation (5 min at 3000 rpm). After addition of 10 uL of 10% SDS, cells were incubated for 10 min at 65°C. SDS was quenched by addition of Triton X-100 and nuclei were resuspended in 300 µL of NEBuffer2 1×. Digestion was performed overnight using 400 U MboI restriction enzyme (NEB). Fill-in of the generated ends with biotinylated-dATP was performed in 300 uL of fresh repair buffer 1× (1.5 µL of 10 mM dCTP, 1.5 µL of 10 mM dGTP, 1.5 µL of 10 mM dTTP, 37.5 µL of 0.4 mM Biotin-dATP, 50 U of DNA Polymerase I large [Klenow] fragment in NEBuffer2 1x) during 45 min at 37°C. Ligation was performed using 10,000 cohesive end units of T4 DNA ligase (NEB) in 1.2 mL of ligation buffer (120 µL of 10× T4 DNA ligase buffer, 100 µL of 10% Triton X-100, 12 µL of 10 mg/mL BSA, 963 µL of H2O) during 4 hours at 16°C. After reversion of the cross-link, purified DNA was fragmented to an average size of 300–400 bp using a Bioruptor Pico (Diagenode; eight cycles; 20 sec on and 60 sec off).

### Hi-C downstream analysis

Fastq files have been processed for quality control and filtering using Fastp (Chen et al., 2018) with default options. Filtered reads have been then used as input for HiC-Pro (Servant et al., 2015) to obtain raw and normalized contact maps for each sample. Conversion from HiC-Pro raw matrices format to hic format has been done using the HiC-Pro utility hicpro2juicebox.sh. ICED-Normalized matrixes have been used for subsequent analysis with GENOVA (van der Weide et al., 2021), whether hic-KR nomalized files have been used with plotgardener (Kramer et al., 2022). HicExplorer (Wolff et al., 2018, Wolff et al., 2020) has also been implemented for generation of h5 files used when the *hicAggregateContacts* function was used for aggregation plots.

## Data and Code Availability

The accession numbers for the raw sequencing and mass spectrometry data reported in this paper are NCBI GEO: GSE291980.

---

**Supplementary Figure 1.**
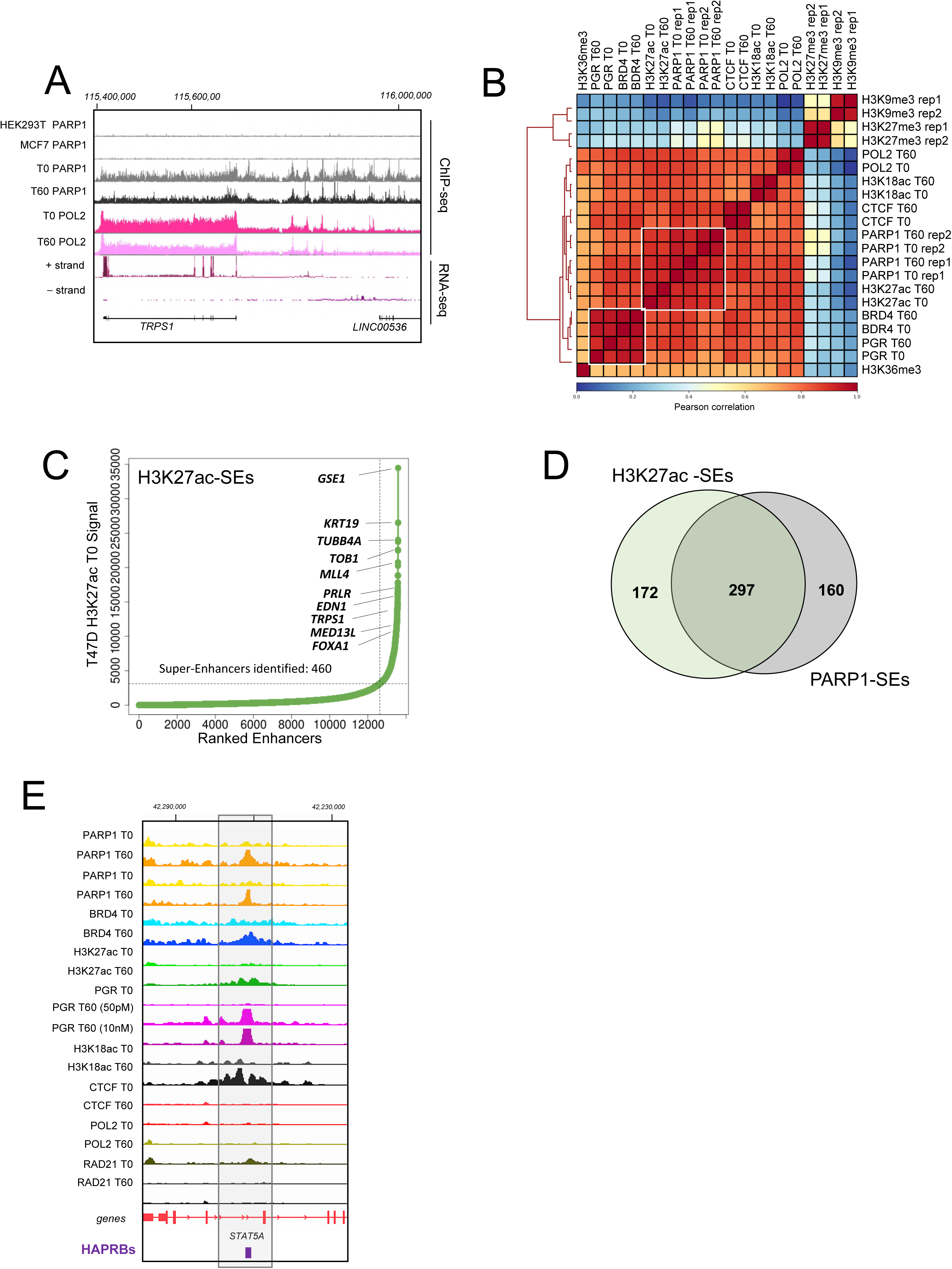
Genome wide PARP1 occupancy correlates with H3K27ac to predict SEs and is recruited to Highly- and Medium-Accessible Progesterone Receptor Binding Sites (HAPRBs, MAPRBs). A) Genome browser view of the representative TRPS1 locus showing successful PARP1 ChIP-seq compared to other PARP1 published ChIP-seq for HEK293T (GSM1938987) and MCF7 (Nalabothula et al., 2015). Note the lack of enrichment for this locus in PARP1 ChIP-seq in HEK293T and MCF7. For this locus are also reported the T47D RNA polymerase II (POL2) ChIP-seq at T0 and T60 and the mRNA-seq at T0. TRPS1 is highly expressed in T47D cells. B) Hierarchical clustering of genome wide Pearson correlations for PARP1 ChIP-seq compared to several histone marks (H3K27ac, H3K18ac, H3K36me3, H3K27me3 and H3K9me3) and chromatin proteins (POL2, BRD4, PGR and CTCF) in T47D. Indicated is the time for the ChIP experiments involving progestin exposure (T0 and T60). PARP1 ChIP-seq replicates clustered close to each other. Notably PARP1 and H3K27ac strongly correlated as highlighted by the white rectangle box. C) Rank ordering of super-enhancers (ROSE) (Loven et al., 2013, Whyte et al., 2013) analysis for H3K27ac ChIP-seq at T0. H3K27ac captures 460 SEs (H3K27ac-SEs) with key T47D genes associated with known SEs from the Super-Enhancer Data Base (SEdb) (Wang et al., 2023). D) Venn diagram reporting the overlap between the PARP1-SEs (Figure 1E) and H3K27ac-SEs. The vast majority of the two sets of SEs overlap. E) Genome browser view of the representative STAT5A locus associated with an HAPRB indicated as a dark purple rectangles). For this locus is shown PARP1 (two replicates), BRD4, H3K27ac, H3K18ac, CTCF, POL2, RAD21 and PGR ChIP-seq in T47D cells at both T0 and T60. For PGR T60 ChIP-seq two different progestin concentrations are shown (50pM and 10nM). Recruitment of PGR to the STAT5A HAPRB increases PARP1 occupancy as well as that of H3K27ac, H3K18ac and BRD4, resulting in increased POL2 recruitment and gene activation as already reported (Zaurin et al., 2021).

**Supplementary Figure 2.**
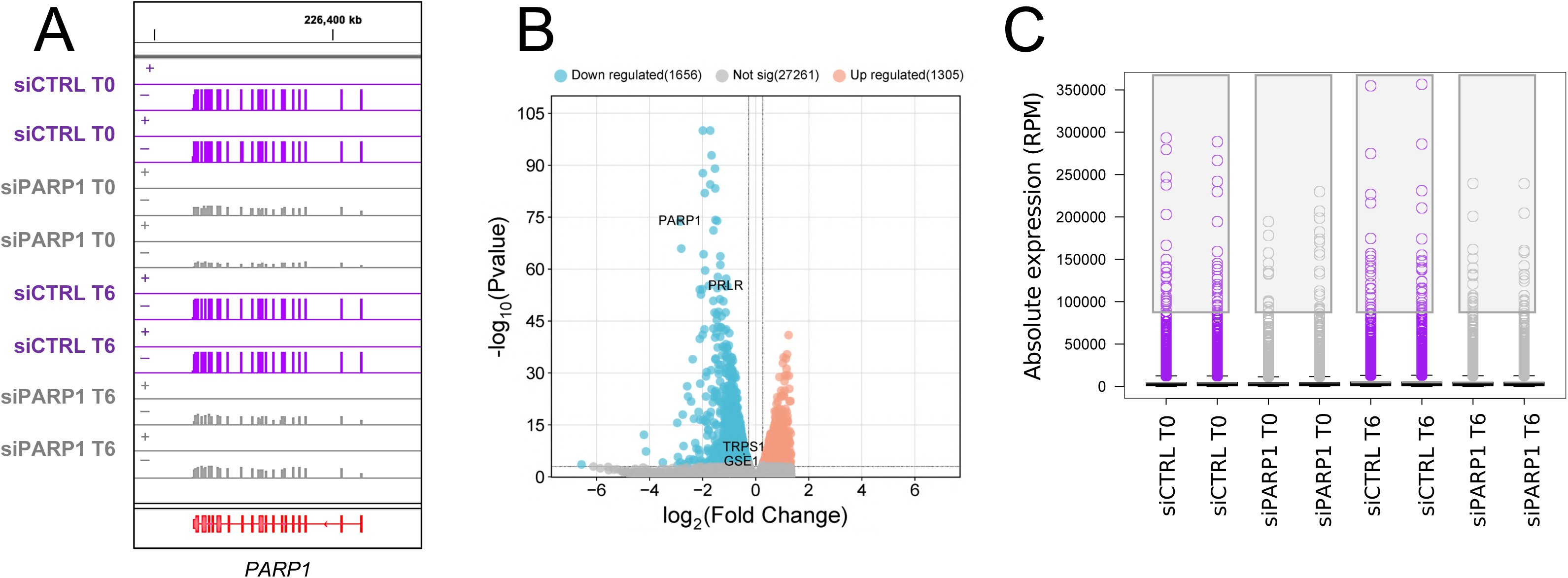
PARP1 KD alters gene expression affecting both SEs and HCRs. **A)** Genome browser view of the PARP1 locus showing the RNA-seq tracks from data of two biological replicates of T47D cells treated with siCTRL T0, siCTRL T60, siPARP1 T0 and siPARP1 T60. At both time points, PARP1 transcript levels strongly decreased. **B)** Volcano plot of gene expression profiling of T47D cells comparing siPARP1 T0 versus siCTRL T0. Indicated are the upregulated genes (red) and downregulated genes (cyan) with respect to a threshold of > 1.5 folds or < −1.5 folds and p-values < 0.05. Most of the genes are downregulated in siPARP1. Indicated are the volcano plot positions for PARP1, PRLR and GSE1 genes. **C)** Boxplot of absolute levels of expression of all genes in each condition indicated. Grey rectangles highlight genes with higher levels of absolute expression that are decreased by the siPARP1 both at T0 and T60. Genes with higher absolute levels of transcription are the most affected by PARP1 KD.

**Supplementary figure 3.**
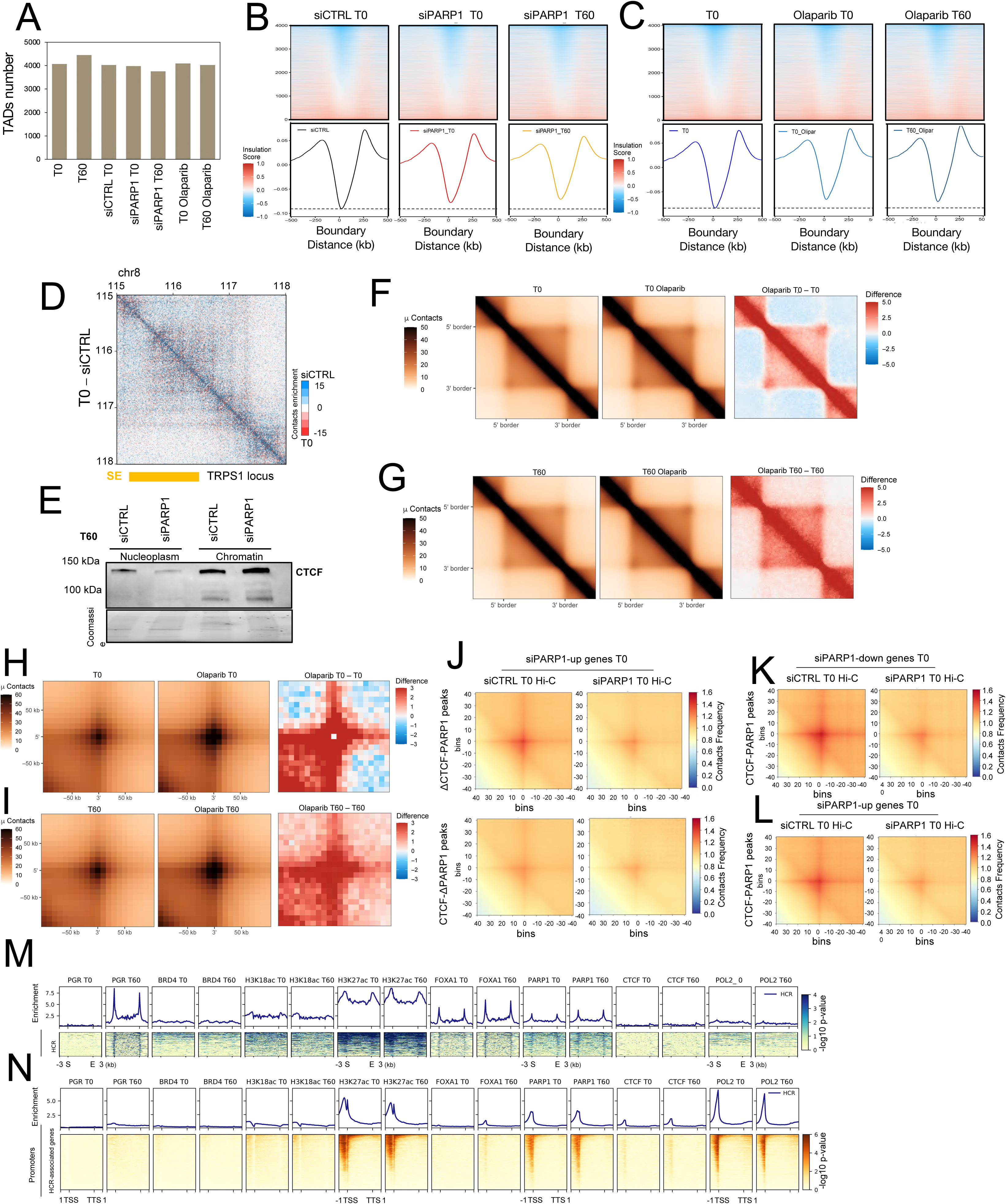
3D genome alteration caused by PARP1 KD. **A)** Computed TADs number for all the Hi-C experiments indicated. No significant variation of TADs number can be observed. TADs have been calculated using hicFindTADs (Ramirez et al., 2018, Wolff et al., 2018, Wolff et al., 2020). **B)** Insulation heatmap and plot across TAD boundaries of siCTRL and siPARP1 at T0, siPARP1 at T60 and **C)** T0 Olaparib treatment at T0 and T60. For this analysis we employed the Insulation-matrix function of GENOVA (van der Weide et al., 2021). **D)** Differential matrix plot for TRPS1 locus comparing Hi-C from siCTRL T0 and T0. SE (indicated at the bottom in orange). Virtual no changes can be detected when using an siCTRL compared to the initial T0 condition. E) Western blot of CTCF upon nuclear fractionation of T47D after R5020 exposure (T60) comparing samples siCTRL and siPARP1. CTCF decreases its level in the nucleoplasm of cells treated with siPARP1 but increases in the chromatin fraction of siPARP1 samples. **F)** Normalized aggregate Hi-C contact heatmaps and differential contact heatmaps around rescaled T0 TADs (n = 4,072) in T0 and T0 Olaparib-treated T47D. **G)** Normalized aggregate Hi-C contact heatmaps and differential contact heatmaps around rescaled T60 TADs (n = 4,466) in T60 and T60 Olaparib-treated T47D. Analysis and visualization of aggregate Hi-C contact heatmaps were performed using GENOVA’s ATA (van der Weide et al., 2021). **H)** Normalized aggregate peak analysis (APA) and differential aggregate peak heatmaps around T0 calculated loops (Wolff et al., 2022), comparing T0 and T0 Olaparib-treated T47D. **I)** Normalized aggregate peak analysis (APA) and differential aggregate peak heatmaps around calculated T60 loops (n = 4,231) comparing T60 and T60 Olaparib-treated T47D. Normalized aggregate peak analysis has been performed using GENOVA’s APA (van der Weide et al., 2021). **J)** Aggregate plots (Ramirez et al., 2018, Wolff et al., 2018, Wolff et al., 2020) of Hi-C interactions between PARP1 peaks devoid of CTCF (ΔCTCF-PARP1, upper figure) and CTCF devoid of PARP1 (ΔCTCF-PARP1, lower figure) with genes up-regulated by PARP1 KD (siPARP1-up genes T0). Compared are the Hi-C data of siCTRL and siPARP1 at T0 for both sets of peaks. **K)** Aggregate plots of Hi-C interactions between PARP1 overlapping CTCF peaks (CTCF-PARP1 peaks) compared to down-regulated genes upon siPARP1 KD at T0 for siCTRL and siPARP1 Hi-C. **L)** Aggregate plots for interactions between PARP1 overlapping CTCF peaks (CTCF-PARP1 peaks) compared to up-regulated genes upon siPARP1 KD at T0 (siPARP1-up genes T0) for siCTRL and siPARP1 Hi-C. **M)** Heatmap showing enrichment for PGR, H3K18ac, H3K27ac, FOXA1, PARP1, CTCF and POL2 (at both T0 and T60) over HCRs. H3K27ac, PARP1 and POL2 show high enrichment. **N)** Heatmap showing enrichment for same factor indicated in panel L over genes affected by R5020 treatment at T60 (Figure 2G). H3K27ac, PARP1 and POL2 show high enrichment.

## Acknowledgments

We acknowledge the support of all members of the Chromatin, Gene Regulation laboratory of Miguel Beato and members of the Gene Regulation Cancer and Stem Cells department at Centre for Genomic Regulation (CRG, Barcelona Spain). R.F. has benefited from the equipment and framework of the COMP-HUB and COMP-R Initiatives, funded by the ‘Departments of Excellence’ Program of the Italian Ministry for University and Research (MIUR, 2018–2022 and MUR, 2022–2027), and from the HPC (High-Performance Computing) facility of the University of Parma, Italy. The experimental part of this manuscript was supported by Grants from ERC (Project “4D Genome” 609989 and Project “Impacct” 825176), the Spanish Ministry of Science (G62426937 and), the Generalitat de Catalunya (AGAUR SGR 757, 2019PROD00115 and INNOV 00036) and the CRG. The experimental part of this manuscript was also supported by Italian Association for Cancer Research [AIRC, IG2022-27712 to R.F.]; Funding for open access charge: AIRC IG2022-27712 to R.F. and R.H.G.W. We acknowledge support of the Spanish Ministry of Science and Innovation to the EMBL partnership the Centro de Excelencia Severo Ochoa and the CERCA Programme/Generalitat de Catalunya.

## Author contributions

Conceptualization, R.H.G.W., R.F. and M.B; writing—original draft preparation, R.H.G.W. and R.F.; writing—review and editing, M.B., R.H.G.W., H.B., F.L. and R.F.; experimental procedures; H.B., R.H.G.W., S.N., A.F.G., F.L and J.F.M. Bioinformatic analysis; R.F., D.C. and J.C.C.; funding acquisition, M.T., M.B., R.H.G.W and R.F. All authors have read and agreed to the published version of the manuscript.

## Inclusion and diversity statement

When your article has been accepted in principle, you will be required to fill out the <u>inclusion and</u> <u>diversity form</u>. In this form, you will have the choice to participate in the initiative by providing pertinent information. If you do provide information, you will then have the option of including this information in an inclusion and diversity statement which will appear in the published paper. You will finally be asked to verify that all authors have agreed to the inclusion of this statement Please note that the information you provide in this form will not have any impact on the decisions we make during consideration of your manuscript. Please find more information on this <u>here</u>, and preview the form <u>here</u>.

